# Genotyping of *Salmonella* spp. on the basis of CRISPR (Clustered Regularly Interspaced Short Palindromic Repeats)

**DOI:** 10.1101/473868

**Authors:** Calvin Jiksing, Normah Yusop, Farhan Nazaie Nasib, Kenneth Francis Rodrigues

**Affiliations:** Biotechnology Research Institute, Universiti Malaysia Sabah, 88400, Kota Kinabalu, Sabah, Malaysia; Diagnostic Veterinary Laboratory, 88999, Kota Kinabalu, Sabah, Malaysia

**Author notes:** Corresponding author: Kenneth Francis Rodriegues, Biotechnology Research Institute, Universiti Malaysia Sabah, 88400, Kota Kinabalu, Sabah, Malaysia, (h/p), 320993. Co-author: Normah Yusop, Diagnostic Veterinary Laboratory, 88999, Kota Kinabalu, Sabah, Malaysia, (h/p); Farhan Nazaie Nasib, Biotechnology Research Institute, Universiti Malaysia Sabah, 88400, Kota Kinabalu, Sabah, Malaysia, (h/p).

**Keywords:** Salmonella, genotyping, serogroup, CRISPR

## Abstract

**Aims:** Bacterial genotyping on the basis of the CRISPR array has been established in *Mycobacterium tuberculosis* with a method called spacer oligonucleotide typing (spoligotyping). The spoligotyping method had been widely used for both detection and typing of *M. tuberculosis* complex bacteria. This present study aimed at determining if the CRISPR array in *Salmonella* spp. could be applied to establish a correlationship between serogroup and the fingerprint generated by CRISPR typing.

**Methodology and results:** A total of 30 samples were obtained from Diagnostic Veterinary Laboratory, Kota Kinabalu, Sabah. Serogroup was determined on the basis of ELISA (enzyme-linked immunosorbent assay). Four different serogroups were identified which were serogroup B, C, D, and E. DNA (deoxyribonucleic acid) was extracted and PCR (polymerase chain reaction) was performed using primers which were designed to amplify the CRISPR array in *Salmonella* genome. Our results indicate that there is a correlationship between serogroup obtained using ELISA and the profile generated by CRISPR typing.

**Conclusion, significance and impact of study:** CRISPR typing has the potential to be applied for the genotyping of *Salmonella*.

## INTRODUCTION

The first CRISPRs locus were identified over 25 years ago in *Escherichia coli* as ambiguous repeat (Ishino *et al*., 1987) and are known as CRISPR spacer arrays now (Mojica *et al*., 2000; Jansen *et al*., 2002a; 2002b). CRISPR arrays consists of tandem direct repeats (DRs) of 23 to 55 bp (base pair) in length interspaced by equal sized of variable spacer sequences that acquired from bacteriophages or plasmids (Bolotin *et al*., 2005; Mojica *et al*., 2005; Pourcel *et al*., 2005; Boyaval *et al*., 2007). The spacer in CRISPR locus was first applied to subtyping *Mycobacterium tuberculosis* strains and this method was known as spacer-oligonucleotide typing or “spoligotyping” (Groenen *et al*., 1993; Kamerbeek *et al*., 1997). Recently, the “next generation” microbead-spoligotyping method was applied to *Salmonella* in an assay named CRISPOL (for “<u>CRIS</u>PR <u>pol</u>ymorphism”)(Fabre *et al*., 2012). Fabre *et al*. (2012) state that there are at least three potential interest of using the polymorphism in *Salmonella* CRISPR locus for clinical microbiology or public health laboratories which are: 1. CRISPR sizing by PCR to compare different isolate of *Salmonella* spp. 2. CRISPOL assay to subtype *Salmonella* serotype Typhimurium or its monophasic variant. 3. Development of PCR assay to target specific *Salmonella* serotype or strain. We aimed to demonstrate that CRISPR based genotyping can be utilized as a diagnostic tool to differentiate different *Salmonella* serogroups prevalent in Sabah.

## MATERIAL AND METHODS

### Sample Collection

*Salmonella* samples (N=30) were obtained from the Diagnostic Veterinary Laboratory, Kota Kinabalu, Sabah. All the *Salmonella* samples were isolated from avian host. The *Salmonella* bacteria were cultured on nutrient agar and MacConkey agar. The MacConkey agar is a selective medium for *Salmonella* which functions to ensure that contaminants are eliminated. The *Salmonella* colony from nutrient agar were transferred to nutrient broth and cultured overnight in incubator shaker at 37°C and 200 rpm (revolutions per minute). The *Salmonella* samples were stored at 25% glycerol stock in −80°C freezer for future use.

### Serogrouping

ELISA was carried out as follows. *Salmonella* were cultured overnight at nutrient agar by streak plate method at 37°C. 0.85% saline solution was used for the ELISA test. The ELISA test was done using the *Salmonella* Sero-Quick Group Kit from Statens Serum Institut, Denmark. The protocol for *Salmonella* ELISA test was done according to the manufacturer protocol.

### DNA Extraction

DNA was extracted according to modified Kang *et al*. (1998) method. The dry DNA pellet was re-suspended in 100 µl of 1X TE (Tris-EDTA) buffer and stored in −20°C for future use.

### PCR of *invA* gene

The primer pair used for this PCR were taken from Cortez *et al*. (2006). PCR was performed in a reaction volume of 25 µl using the GE Healthcare illustra^TM^ puReTaq Ready-To-Go PCR Beads. The preparation of PCR master mix was prepared according to manufacturer protocol. Amplification was carried out in a thermal cycler (MJ Research PTC-200 Peltier Thermal Cycler) using 35 cycles consisting of denaturation for 30 sec at 94°C, annealing for 1 min at 55°C, and extension for 1 min at 72°C, followed by a final extension for 7 min at 72°C. Electrophoresis of amplified products was carried out using 1.0% agarose gel in 1X TBE (Tris/Borate/EDTA) running buffer. The amplified DNA fragments were stained with ethidium bromide and visualized under UV (ultraviolet) light. A 100 bp DNA ladder (New England Biolabs Quick-Load 100 bp DNA Ladder) was used as a reference standard.

### Primer Design

Design of CRISPR specific primers was carried out as follows. The complete genome of *Salmonella enterica* subsp. *enterica* serovar Typhi strain CT18 was first retrieved from the NCBI (National Centre for Biotechnology Information) with accession number of NC_003198.1. The genome was then analyzed for presence of CRISPR locus using online web tool called CRISPERFinder at http://crispr.upsud.fr/Server/CRISPRfinder.php. Once the CRISPR locus sequence was identified three pairs of primer set were designs to amplify the CRISPR 1 and CRISPR 2 locus presence in *Salmonella* genome using online web tool called Primer3 at http://bioinfo.ut.ee/primer3-0.4.0/. The primer sequences are listed in Table 1.0.

**Table 1.0.**
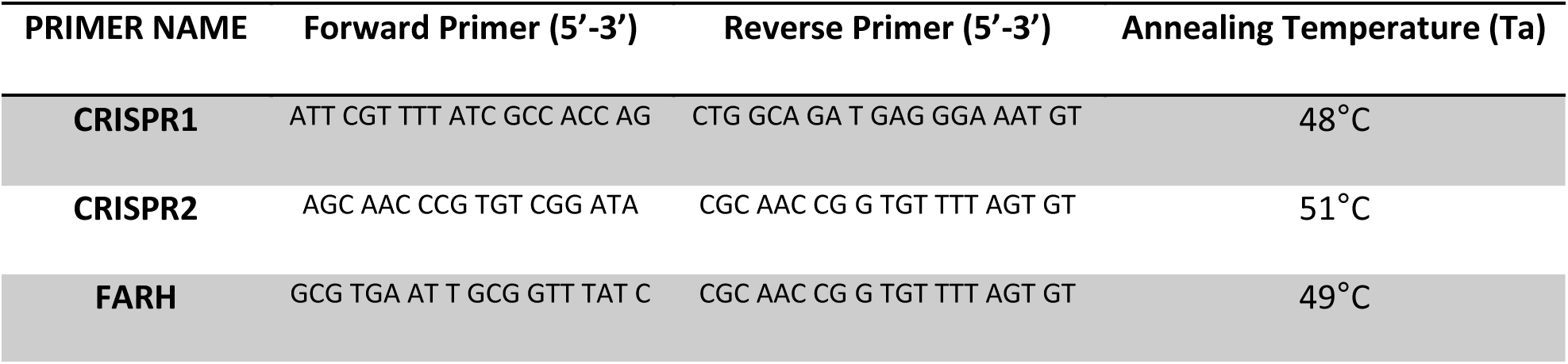
List of primer used for the CRISPR typing.

### PCR of CRISPR Locus

PCR was performed in a reaction volume of 20 µl containing 1X Dream Taq. Green Buffer (Thermo Scientific), 0.1 mM dNTPs (deoxynucleoside triphosphate), 2 mM MgCl_2_ (Magnesium chloride), 5 µM of forward and reverse primer, 1 Unit of Thermo Scientific Dream Taq DNA polymerase and 1 µl of DNA template. Amplification was carried out in a thermal cycler (MJ Research PTC-200 Peltier Thermal Cycler) with initial denaturation of 96°C for 4 min, followed by 35 cycles of 96°C for 30 sec, 51°C for 30 sec, 72°C for 1 min and final extension step at 72°C for 2 min. All steps were the same for all primers except for the annealing temperature. The annealing temperature for CRISPR1 primer pair is 48°C, 51°C for CRISPR2 primer pair and 49°C for FARH primer pair. Electrophoresis of amplified products was carried out using 1.0% agarose gel in 1X TBE running buffer. The amplified DNA fragments were stained with ethidium bromide and visualized under UV light. A 100 bp DNA ladder (New England Biolabs Quick-Load 100 bp DNA Ladder) was used as a reference standard.

## RESULTS

*Salmonella* samples (N=30) were test by ELISA for serogrouping. 53% of the samples were from serogroup C, followed by serogroup E with 20%, 17% for serogroup B and 10% for serogroup D. The genus confirmations for all the samples were done by the PCR of *invA* gene (Figure 1.0, 2.0 and 3.0). The expected size for the PCR product is 521 bp. PCR to amplify the CRISPR locus on *Salmonella* genome were done to differentiate the different *Salmonella* serogroups (Figure 4.0, 5.0 and 6.0).

**Figure 1.0.**
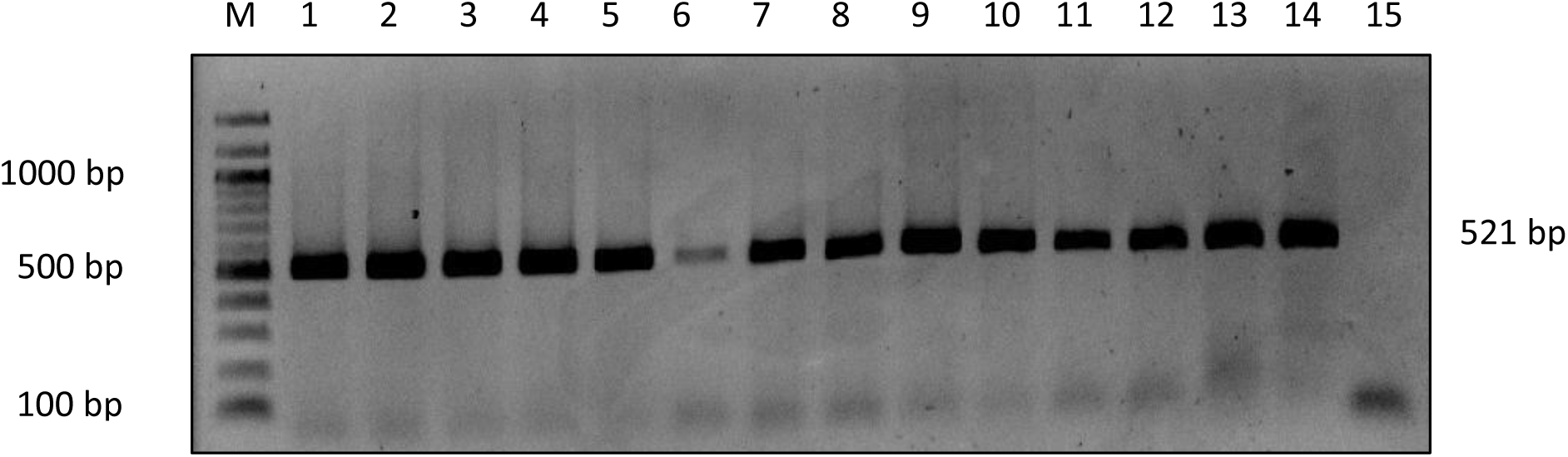
PCR was done to amplify the *invA* gene on the *Salmonella* bacteria genome for genus confirmation. Lane M: 100 bp marker, lane 1 to 14: The PCR product of *Salmonella invA* gene and lane 15: Negative control without DNA template.

**Figure 2.0.**
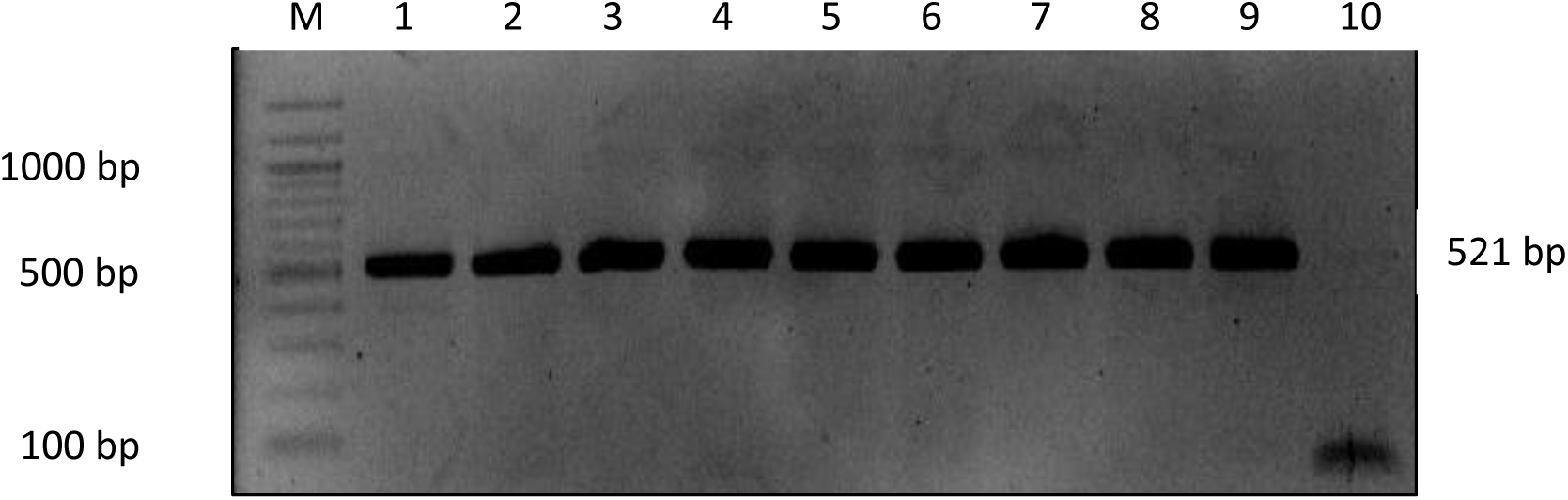
PCR was done to amplify the *invA* gene on the *Salmonella* bacteria genome for genus confirmation. Lane M: 100 bp marker, lane 1 to 9: The PCR product of *Salmonella invA* gene and lane 10: Negative control without DNA template.

**Figure 3.0.**
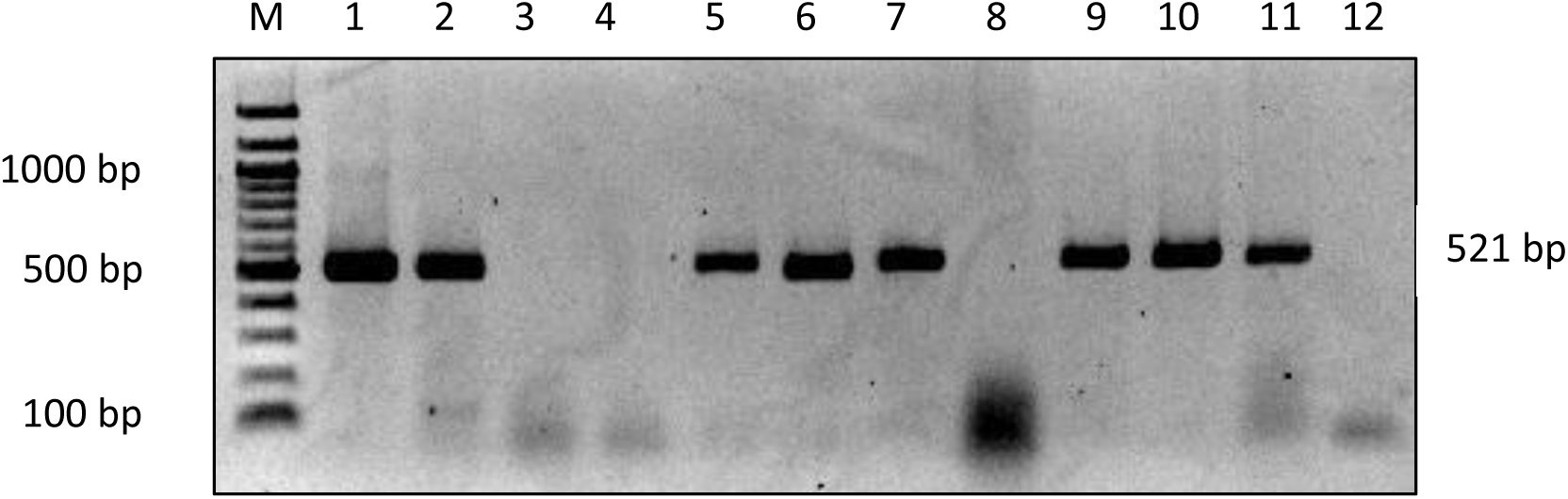
PCR was done to amplify the *invA* gene on the *Salmonella* bacteria genome for genus confirmation. Lane M: 100 bp marker, lane 1 to 11: The PCR product of *Salmonella invA* gene and lane 12: Negative control without DNA template.

**Figure 4.0.**
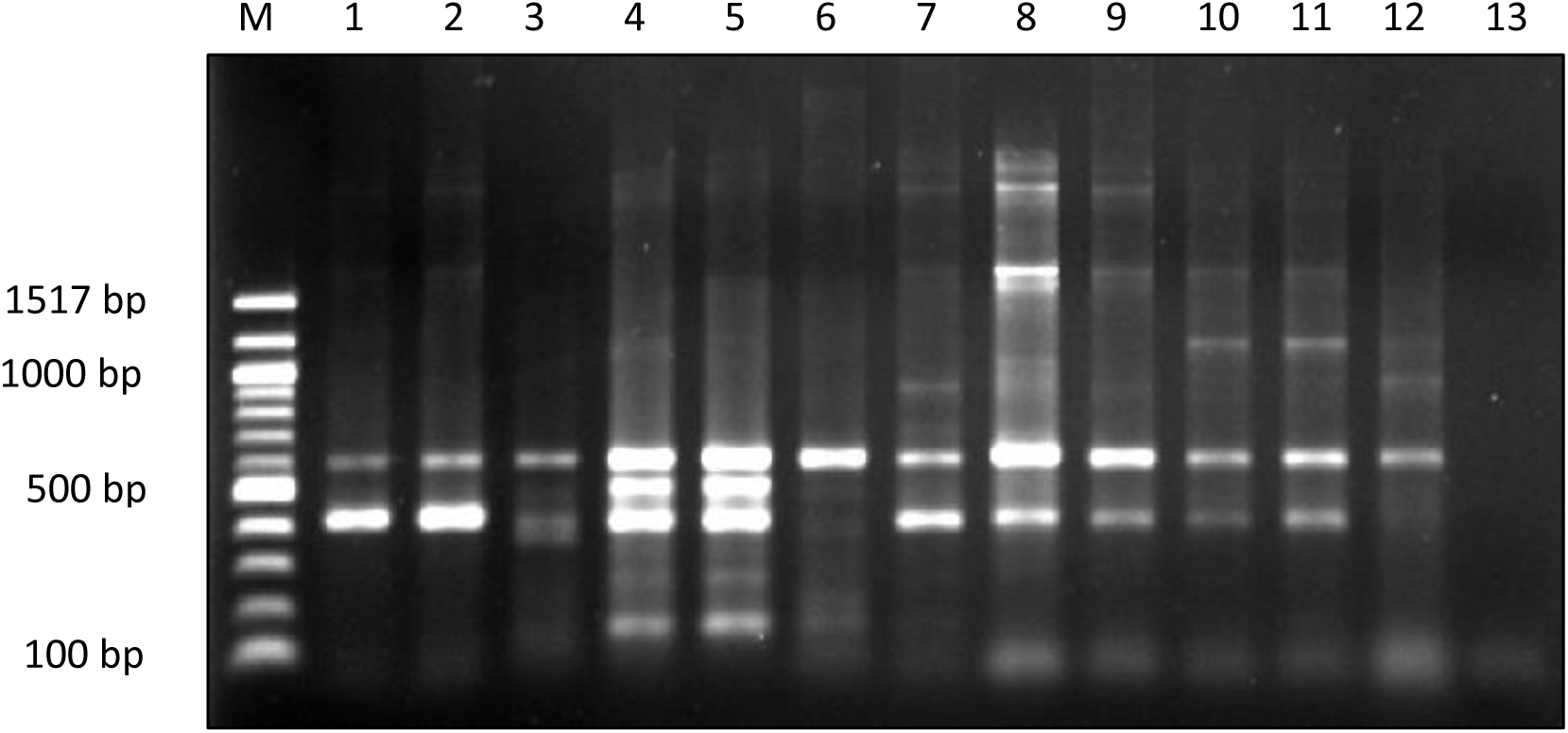
CRISPR1 primer pair was designed for PCR to amplify the CRISPR locus in *Salmonella* genome for serogrouping differentiation by molecular method. The above agarose gel picture show the amplification profile when *Salmonella* genome was tested with the designed primer. Lane M: 100 bp marker, lane1-3: *Salmonella* from serogroup B, lane 4-6: *Salmonella* from serogroup C, lane 7-9: *Salmonella* from serogroup D, lane 10-12: *Salmonella* from serogroup E and lane 13: Negative control without DNA template.

**Figure 5.0.**
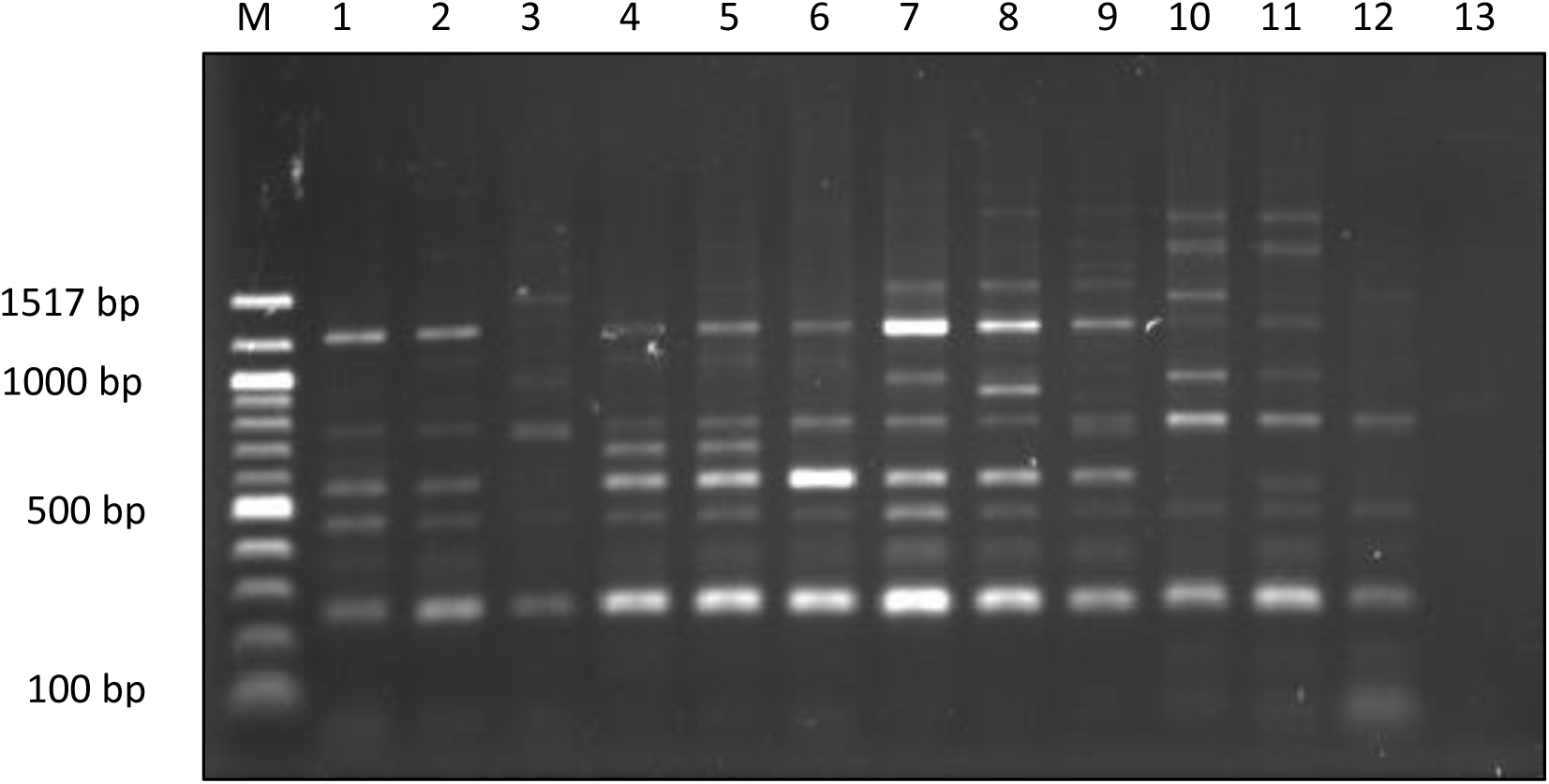
CRISPR2 primer pair was designed for PCR to amplify the CRISPR locus in *Salmonella* genome for serogrouping differentiation by molecular method. The above agarose gel picture show the amplification profile when *Salmonella* genome was tested with the designed primer. Lane M: 100 bp marker, lane1-3: *Salmonella* from serogroup B, lane 4-6: *Salmonella* from serogroup C, lane 7-9: *Salmonella* from serogroup D, lane 10-12: *Salmonella* from serogroup E and lane 13: Negative control without DNA template.

**Figure 6.0.**
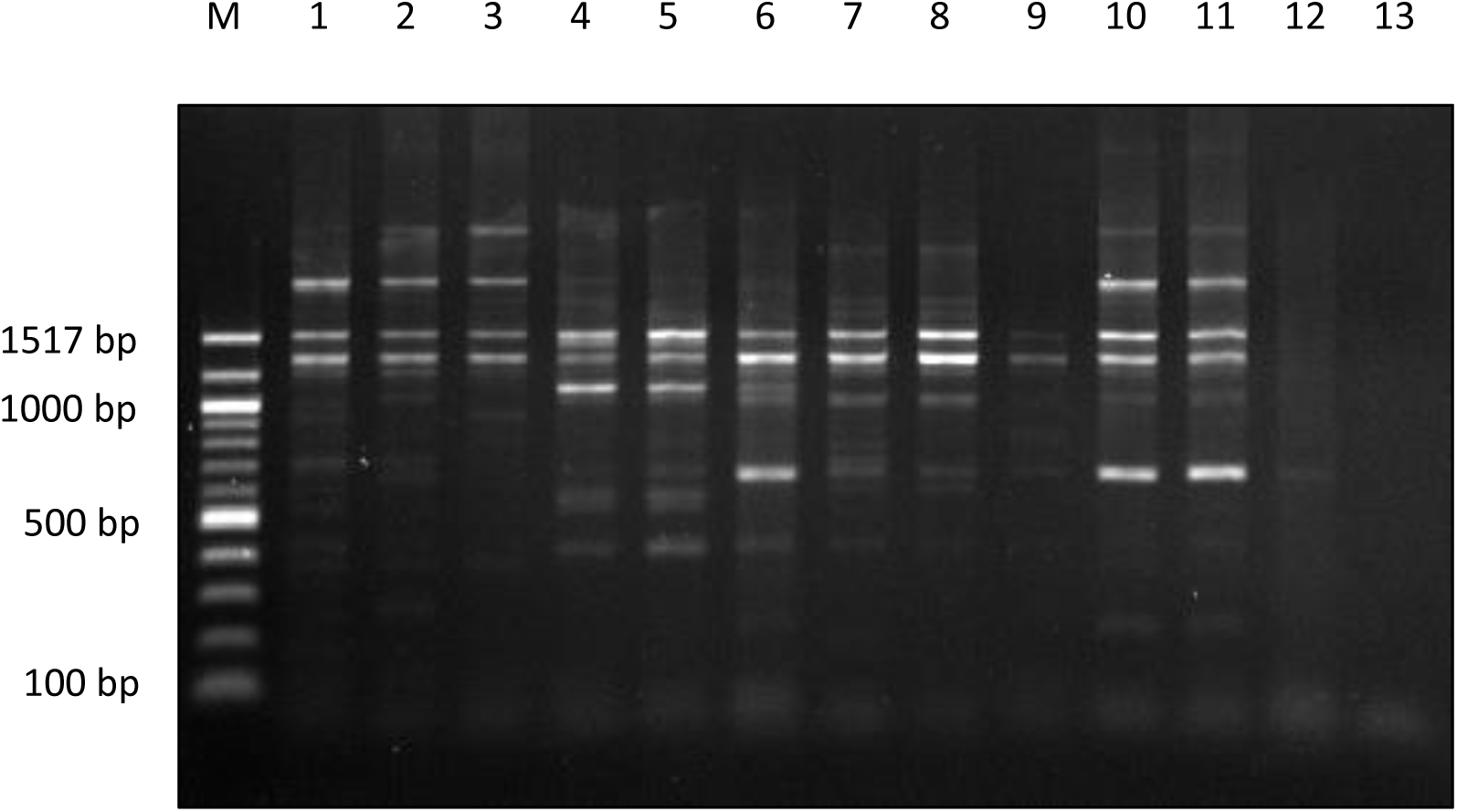
FARH primer pair was designed for PCR to amplify the CRISPR locus in *Salmonella* genome for serogrouping differentiation by molecular method. The above agarose gel picture show the amplification profile when *Salmonella* genome was tested with the designed primer. Lane M: 100 bp marker, lane1-3: *Salmonella* from serogroup B, lane 4-6: *Salmonella* from serogroup C, lane 7-9: *Salmonella* from serogroup D, lane 10-12: *Salmonella* from serogroup E and lane 13: Negative control without DNA template.

## DISCUSSION

The O antigen present in the cell surface of *Salmonella* is extremely polymorphic and is used to determine the bacteria serogroup (Bee & Kwai, 2009). The variation in O antigen structure is due to the different types of sugar present, the arrangement of sugars, the addition of branch sugars and the modifying side groups in which such variation is used to serogroup *Salmonella* isolates (Wyk & Reeves, 1989; Fitzgerald *et al*., 2003; Luk *et al*., 2006). In this study four serogroups were found which were serogroup B, C, D and E. This finding is similar with the finding of Lindberg & Le Minor (1984) and Luk & Lindberg (1991) that stated over than 95% of the *Salmonella* strains that cause infection in human and animal is originate from the serogroup A to E. The identity of the *Salmonella* samples recruited for this study was confirmed by the present of *invA* gene in *Salmonella* genome using PCR method. This was done in order to ensure that all the samples used in this study were from the pure culture of *Salmonella* bacteria. PCR was an effective, rapid, reliable and sensitive method for the detection of *invA* gene present in *Salmonella* genome (Zahraei-Salehi *et al*., 2006). A study from Galán & Curtiss (1989) show that a group of genes (*invA, B, C, D*) confer *Salmonella* the ability to invade cultured epithelial cells. The use of this gene for *Salmonella* identification by PCR method has recently been suggested as this gene were shown to be found in a number of *Salmonella* strain (Galán & Curtiss, 1991). A study by Zahraei-Salehi *et al*. (2006) confirm that the *invA* gene sequence is unique to *Salmonella* and this unique sequence can be used as the PCR target to differentiate *Salmonella* from other organisms. The CRISPR typing was done to differentiate *Salmonella* serogroup by molecular method and the result was compared with the traditional serogrouping by ELISA method. The amplification profile of *Salmonella* from different serogroups were observed and compared. The principle of this method is that the variation in spacer number that interspaced between the direct repeats will give different lengths of CRISPR array and can be used to rapidly screen *Salmonella* isolates by PCR and gel electrophoresis analysis (Shariat & Dudley, 2014). The CRISPR1 and CRISPR2 primer pairs failed to differentiate some of the samples when compared to the serogrouping result obtained by the ELISA method. Depending on the antibody use for ELISA test, they may lack specificity because of the non-specific agglutination that might happen in some *Salmonella* bacteria as stated by Cheesbrough & Donnelly (1996). Only the CRISPR primer designated as FARH primer was able to completely differentiate all the samples from different serogroups. The reason why CRISPR1 and CRISPR2 primer cannot resolve the serogroup while FARH primer can might be because of the location of the primer along the CRISPR gene. The positions of forward and reverse primer for CRISPR1 along the gene are at nucleotide number 2,925,620 to 2,925,640 and 2,926,778 to 2,926,798 respectively. For CRISPR2 the positions of the forward and reverse primer along the gene are at nucleotide number 2,926,215 to 2,926,233 and 2,926,450 to 2,926,470 respectively. For the FARH the positions of forward and reverse primer along the gene are at nucleotide number 2,926,294 to 2,926,313 and 2,926,450 to 2,926,470 respectively. It can be seen from the results that some samples from the same serogroup have different amplification profile. This might be because of in one serogroup there were sub-group present as shown in White-Kauffmann-Le-Minor scheme (Grimont & Weill, 2008). Our findings are consistent with those obtained by Fabre *et al*. (2012) and Shariat & Dudley (2014) who used rapid CRISPR size typing to screen *Salmonella* spp. isolates by comparing the amplicon size of PCR product run on agarose gel electrophoresis.

## CONCLUSION

From this study, it shown that only the FARH primer is able to resolve the samples into their respective serogroup. The serogrouping by ELISA method can be complement with CRISPR typing by PCR method to get more accurate result as non-specific agglutination might occur using the ELISA method.

## ACKNOWLEDGEMENTS

We would like to thank UMS (Universiti Malaysia Sabah) for supporting this project under the UMSGreat grant with code project of GUG0106-1/2017. We also would like to thank the Veterinary Laboratory, Kota Kinabalu, Sabah for providing us the *Salmonella* samples used in this study.

## REFERENCES

Bee, K. L. and Kwai, L. T. (2009). Application of PCR-based serogrouping of selected Salmonella serotypes in Malaysia. Journal of Infection in Developing Countries 3(6), 420–428.

Bolotin, A., Quinquis, B., Sorokin, A. and Dusko Ehrlich, S. (2005). Clustered regularly interspaced short palindrome repeats (CRISPRs) have spacers of extrachromosomal origin. Microbiology 151(8), 2551–2561.

Boyaval, P., Moineau, S., Romero, D. A. and Horvath, P. (2007). Against Viruses in Prokaryotes. Science 315, 1709–1712.

Cheesbrough, S. and Donnelly, C. (1996). The use of a rapid Salmonella latex serogrouping test (Spectate(®)) to assist in the confirmation of ELISA-based rapid Salmonella screening tests. Letters in Applied Microbiology 22(5), 378–380.

Cortez, A. L. L., Carvalho, A. C. F. B., Ikuno, A. A., Bürger, K. P. and Vidal-Martins, A. M. C. (2006). Identification of Salmonella spp. isolates from chicken abattoirs by multiplex-PCR. Research in Veterinary Science 81(3), 340–344.

Fabre, L., Zhang, J., Guigon, G., Le Hello, S., Guibert, V., Accou-Demartin, M., de Romans, S., Lim, C., Roux, C., Passet, V., Diancourt, L., Guibourdenche, M., Issenhuth-Jeanjean, S., Achtman, M., Brisse, S., Sola, C. and Weill, F.X. (2012). Crispr typing and subtyping for improved Laboratory surveillance of Salmonella infections. PLoS ONE 7(5), e36995.

Fitzgerald, C., Sherwood, R., Gheesling, L. L., Brenner, F. W. and Fields, P. I. (2003). Molecular Analysis of the rfb O Antigen Gene Cluster of Salmonella enterica Serogroup O : 6 , 14 and Development of a Serogroup-Specific PCR Assay. Applied and Environmental Microbiology 69(10), 6099–6105.

Galán, J. E. and Curtiss, R. (1989). Cloning and molecular characterization of genes whose products allow Salmonella Typhimurium to penetrate tissue culture cells. Proc. Natl. Acad. Sci. USA 86, 6383–6387.

Galán, J. E. and Curtiss, R. (1991). Distribution of the invA, -B, -C, and -D genes of Salmonella Typhimurium among other Salmonella serovars: invA mutants of Salmonella Typhi are deficient for entry into mammalian cells. Infection and Immunity 59(9), 2901–2908.

Grimont, P. and Weill, F. X. (2008). Antigenic formulae of the Salmonella serovars. WHO Collaborating Centre for Reference and Research on Salmonella, 1–167. Retrieved from http://www.pasteur.fr/ip/portal/action/WebdriveActionEvent/oid/01s-000036-089%5Cnpapers2//publication/uuid/CA3447A0-61BF-4D62-9181-C9BA78AF0312

Groenen, P. M. A., Bunschoten, A. E., Soolingen, D. V. and Errtbden, J. D. A. V. (1993). Nature of DNA polymorphism in the direct repeat cluster of Mycobacterium tuberculosis; application for strain differentiation by a novel typing method. Molecular Microbiology 10(5), 1057–1065.

Ishino, Y., Shinagawa, H., Makino, K., Amemura, M. and Nakata, A. (1987). Nucleotide sequence of the iap gene, responsible for alkaline phosphatase isozyme conversion in Escherichia coli, and identification of the gene product. Journal of Bacteriology 169(12), 5429–5433.

Jansen, R., van Embden, J. D. A., Gaastra, W. and Schouls, L. M. (2002a). Identification of genes that are associated with DNA repeats in prokaryotes. Molecular Microbiology 43(6), 1565–1575.

Jansen, R., van Embden, J. D. A., Gaastra, W. and Schouls, L. M. (2002b). Identification of a novel family of sequence repeats among prokaryotes. Omics 6(1), 23–33.

Kamerbeek, J., Schouls, L., Kolk, A., Agterveld, M. V., Soolingen, D.V., Kuijper, S., Bunschoten, A., Molhuizen, H., Shaw, R., Goyal, M. and Embden, J.V. (1997). Simultaneous detection and strain differentiation of Mycobacterium tuberculosis for diagnosis and epidemiology Journal of Clinical Microbiology 35(4), 907–914.

Kang, H. W., Cho, Y. G., Yoon, U. H. and Eun, M. Y. (1998). A rapid DNA extraction method for RFLP and PCR analysis from a single dry seed. Plant Molecular Biology Reporter 16, 1–9.

Lindberg, A. A. and Le Minor, L. (1984). Serology of Salmonella. Methods in Microbiology 15, 1–141.

Luk, J. M. C., Kongmuang, U., Reeves, P. R. and Lindberg, A. A. (2006). Selective amplification of abequose and paratose synthase genes (rfb) by polymerase chain reaction for identification of Salmonella major serogroups (A, B, C2, and D). Journal of Clinical Microbiology 31(8), 2118–2123.

Luk, J. M. C. and Lindberg, A. A. (1991). Anti-Salmonella lipopolysaccharide monoclonal antibodies: Characterization of Salmonella BO-, CO-, DO-, and EO-specific clones and their diagnostic usefulness. Journal of Clinical Microbiology 29(11), 2424–2433.

Mojica, F. J. M., Díez-Villaseñor, C., García-Martínez, J. and Soria, E. (2005). Intervening sequences of regularly spaced prokaryotic repeats derive from foreign genetic elements. Journal of Molecular Evolution 60(2), 174–182.

Mojica, F. J. M., Díez-Villaseñor, C., Soria, E. and Juez, G. (2000). Biological significance of a family of regularly spaced repeats in the genomes of Archaea, Bacteria and mitochondria. Molecular Microbiology 36(1), 244–246.

Pourcel, C., Salvignol, G. and Vergnaud, G. (2005). CRISPR elements in Yersinia pestis acquire new repeats by preferential uptake of bacteriophage DNA, and provide additional tools for evolutionary studies. Microbiology 151(3), 653–663.

Shariat, N. and Dudley, E. G. (2014). CRISPRs : Molecular Signatures Used for Pathogen Subtyping. Applied and Environmental Microbiology 80(2), 430–439.

Wyk, P. and Reeves, P. (1989). Identification and sequence of the gene for abequose synthase, which confers antigenic specificity on group B salmonellae: Homology with galactose epimerase. Journal of Bacteriology 171(10), 5687–5693.

Zahraei-Salehi, M.T., Mahzoniae, M. R. and Ashrafi, A. (2006). Amplification of invA gene of Salmonella by polymerase chain reaction (PCR) as a specific method for detection of Salmonellae. Journal of Veterinary Research 61(2), 195–199.

